# Distinct cAMP regulation in scleroderma lung and skin myofibroblasts governs their dedifferentiation via p38α inhibition

**DOI:** 10.1101/2025.02.26.640163

**Authors:** Jared D. Baas, John Varga, Carol Feghali-Bostwick, Marc Peters-Golden, Sean M. Fortier

**Affiliations:** Division of Pulmonary and Critical Care Medicine, University of Michigan Medical School, Ann Arbor, Michigan, USA; Division of Rheumatology, University of Michigan Medical School, Ann Arbor, Michigan, USA; Division of Rheumatology, Medical University of South Carolina, Charleston, USA

## Abstract

Fibrosis in systemic sclerosis/scleroderma (SSc) is characterized by the progressive accumulation and persistence in multiple organs of pathologic fibroblasts whose contractile properties and exuberant secretion of collagens promote tissue stiffness and scarring. Identifying a tractable mechanism for inactivating and possibly clearing these ultimate effector cells of fibrosis, conventionally termed myofibroblasts (MFs), represents an appealing therapeutic strategy for patients with SSc. This can be accomplished by their phenotypic dedifferentiation, a process known to be promoted by generation of the intracellular second messenger cyclic AMP (cAMP). Notably, however, the abilities of SSc fibroblasts derived from different tissues to generate cAMP – and dedifferentiate in response to it – have never been directly characterized or compared. Here we compared these two processes in lung and skin MFs derived from patients with SSc. While directly increasing intracellular cAMP induced comparable dedifferentiation of lung and skin SSc MFs, dedifferentiation in response to the well-recognized cAMP stimulus prostaglandin E_2_ (PGE_2_) was diminished or absent in MFs from skin as compared to lung, in part due to differences in the expression of its target G protein-coupled receptors (GPCRs). Importantly, treatment with a phosphodiesterase 4 inhibitor rescued the dedifferentiating effects of PGE_2_ in skin SSc MFs. Finally, both cAMP-mediated and direct pharmacologic inhibition of the MAPK p38α promoted dedifferentiation of lung and skin SSc MFs. We conclude that activation of the cAMP pathway and its subsequent inhibition of p38α dedifferentiates SSc MFs from both lung and skin, and may thus represent a therapeutic strategy to alleviate multi-organ fibrosis in SSc.

## INTRODUCTION

Fibrosis is the final common consequence of repetitive tissue injury and a primary etiology of organ dysfunction and mortality in numerous chronic disorders (1). Systemic sclerosis (SSc) is an autoimmune connective tissue disease characterized by chronic vascular injury, inflammation, and fibrosis of the skin and internal organs (2). The ultimate effector cell responsible for organ stiffness and scarring in SSc is the myofibroblast (MF) – a pathologically activated mesenchymal cell derived mainly from the differentiation of quiescent resident tissue fibroblasts that produces excess collagens/matrix proteins, contributing to contractile forces that distort tissue architecture and disrupt its functions (3–5).

Persistence and accumulation of MFs distinguish SSc and other fibrotic disorders from normal wound healing – in which their timely clearance is associated with restoration of tissue structure and homeostasis (6). Indeed, lung MF clearance in murine models of spontaneous fibrosis resolution requires their phenotypic dedifferentiation (7, 8) – marked by a decline in fibrosis-associated proteins (αSMA and collagen I) together with a dismantling of intracellular stress fibers – and eventual reacquisition of their capacity for apoptosis (9). Despite the potential therapeutic importance of reversing the MF phenotype via dedifferentiation, the overwhelming preponderance of studies involving tissue fibroblasts have instead focused on merely preventing their differentiation into MFs in response to profibrotic stimuli such as transforming growth factor β-1 (TGFβ). Yet, treatments that promote and potentiate MF dedifferentiation are more likely to be clinically impactful, particularly as patients with chronic fibrotic disorders such as SSc often exhibit established organ fibrosis by the time they reach clinical attention.

It is now well established that tissue MFs can be dedifferentiated *in vitro* by treatment with a variety of mediators and pharmacologic agents (10–13). Among the most active and best-studied dedifferentiating agents are those that signal by increasing intracellular levels of the second messenger cyclic AMP (cAMP), which is also known to inhibit all major functions within fibroblasts, including proliferation, collagen synthesis, differentiation, and migration (14–18). Intracellular cAMP concentration depends on the balance of its a) generation – in response to engagement of stimulatory (Gαs) or inhibitory (Gαi) G-protein coupled receptors (GPCRs) that activate or inhibit, respectively, adenylyl cyclase (AC) – and b) degradation via phosphodiesterases (PDEs) that catalyze the conversion and degradation of cAMP into AMP. Importantly, the nature and capacity of the cellular response to mediators that signal via cAMP varies by cell type, tissue, and states of disease. For instance, fibroblasts derived from some patients with idiopathic pulmonary fibrosis (IPF) exhibit an impaired capacity to increase intracellular cAMP (19).

Current therapeutic options for SSc primarily target immune cell function (20) and fail to modulate the mesenchymal cells directly responsible for scarring. We therefore sought to determine and characterize the effect of increasing intracellular cAMP within SSc patient-derived MFs. SSc is unique among fibrotic conditions in that patients exhibit fibrosis within multiple organs, highlighting the biological importance of determining common and divergent features and responses among tissue resident fibroblasts (21). Skin fibrosis is an early manifestiation of SSc and a major source of morbidity (22), while SSc-related lung fibrosis (SSc-ILD) confers the highest risk for mortality (23). We therefore characterized and compared the ability of MFs obtained from the lungs and skin of patients with SSc to undergo dedifferentiation in response to direct increases in intracellular cAMP, as well as their capacity to generate cAMP and undergo dedifferentiation in response to PGE_2_, a canonical endogenous mediator coupled to cAMP production. As cAMP is known to influence cell phenotype in part via MAPK proteins (7), its effect on downstream p38, Erk, and JNK activity was also assessed.

We found that generation of intracellular cAMP via direct activation of AC promoted a comparable degree of dedifferentiation in MFs from both SSc lung and skin, whereas PGE_2_ exclusively dedifferentiated lung. This lack of effect in skin MFs reflected impaired capacity of PGE_2_ to increase intracellular cAMP. Importantly, the inability of PGE_2_ to increase cAMP and elicit dedifferentiation in skin MFs was overcome by addition of a PDE4 inhibitor. Moreover, PDE4 inhibition alone reduced αSMA expression and dismantled stress fibers in both lung and skin MFs. Finally, we linked the dedifferentiating actions of cAMP to p38 inhibition – specifically the alpha isoform – in SSc MFs from both lung and skin. Our results highlight an important organ-specific difference in PGE_2_-mediated cAMP production in SSc MFs while demonstrating the common dedifferentiating effects of cAMP, the substantial effect of PDE4 activity on cAMP levels within SSc skin MFs, and the cAMP-mediated inhibition of p38α within SSc fibroblasts from both organs. These results provide novel insights into potential mechanisms by which the growing armamentarium of agents that engage the cAMP pathway regulate mesenchymal cells from various tissues.

## RESULTS

### Directly increasing intracellular cAMP promotes dedifferentiation in both lung and skin MFs from SSc patients

Pilot studies comparing gene expression in lung and skin fibroblasts derived from healthy individuals to those from patients with SSc revealed a similar degree of baseline, and TGFβ-upregulated, fibrosis-associated transcripts in culture (data not shown). We therefore chose to focus our investigation on SSc patient-derived cells. To evaluate the capacity of cAMP to dedifferentiate established SSc lung and skin MFs, we first treated SSc fibroblasts with TGFβ for 48 h to unequivocally induce the MF phenotype (Figure 1A). Next, SSc lung and skin MFs were treated with the direct adenylyl cyclase (AC) activator, forskolin, or the cell-permeable cAMP analog dibutyryl cAMP (db-cAMP), the actions of which are depicted in the Figure 1B schematic. We previously reported that in normal lung fibroblasts, cAMP signaling in response to PGE_2_ results in the downregulation of a broad array of genes in fibrosis-associated pathways including those associated with the extracellular matrix, cytoskeletal adhesion, and focal adhesion, as well as TGFβ, Wnt, mTOR, calcium, and PI3 kinase signaling (7). Here we employed the representative canonical fibrosis-associated markers αSMA and Col1A1 as readouts of the MF phenotype. Both forskolin and db-cAMP robustly decreased these markers at both the protein (Figure 1C) and transcript (Supplemental Figure 1A and 1B) level within established SSc lung and skin MFs. Furthermore, immunofluorescence microscopy revealed that both drugs substantially reduced the organization of αSMA into stress fibers and elicited a conspicuous morphologic transition – thinner cells with decreased cytoplasm – typical of phenotypic dedifferentiation (Figure 1D). These results indicate that direct increases in intracellular cAMP promote the dedifferentiation of both lung and skin MFs derived from patients with SSc.

**Figure 1.**
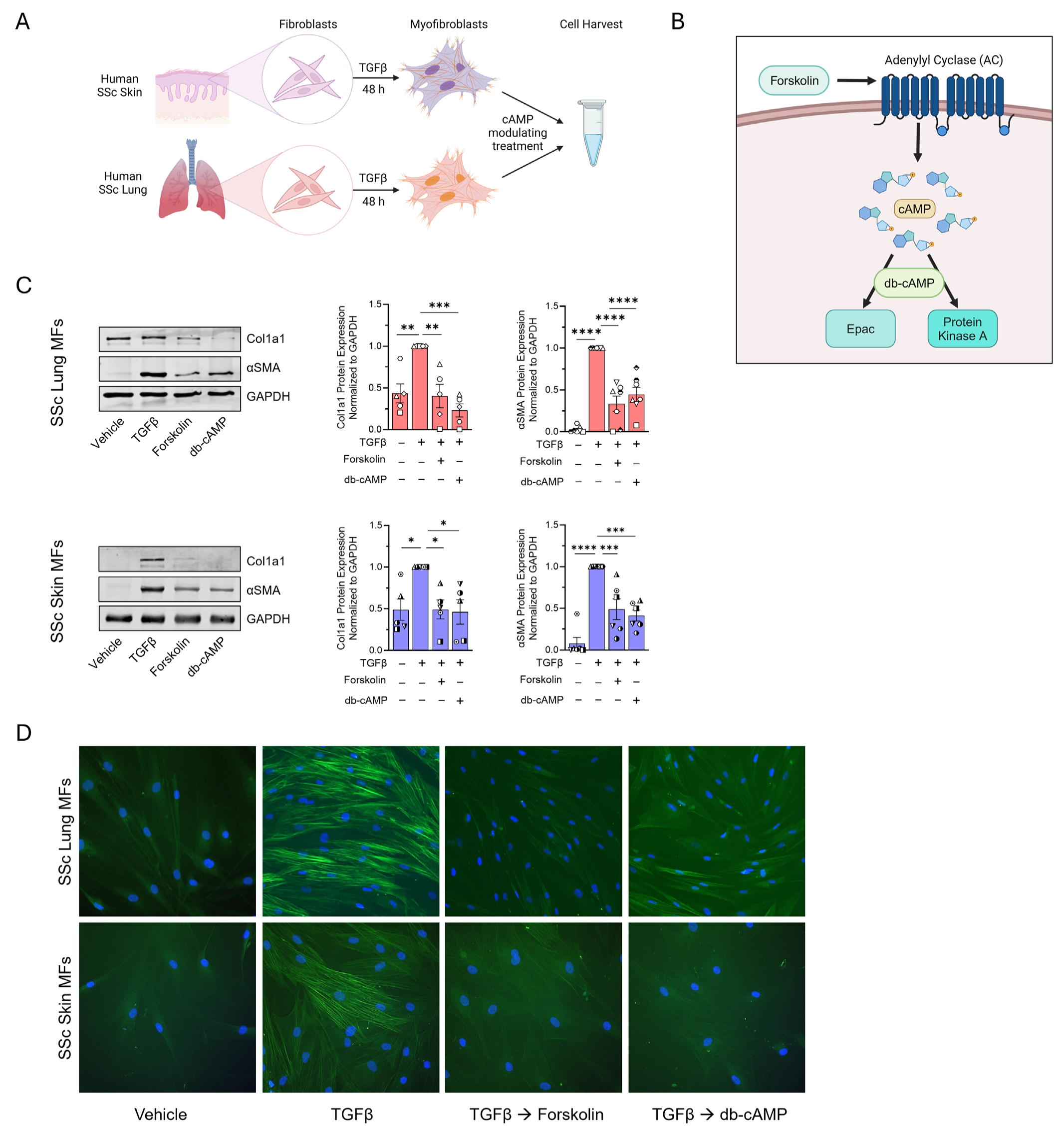
Directly increasing intracellular cAMP promotes dedifferentiation in both lung and skin MFs from SSc patients. (**A**) Schematic depicting the MF differentiation and dedifferentiation protocols employed. SSc fibroblasts derived from skin or lungs were treated with TGFβ (2 ng/mL) for 48 h, followed by administration of a cAMP modulating treatment for 48 to 96 h before harvest and measurement of mRNA or protein, respectively. (**B**) Illustration demonstrating the experimental actions of forskolin and db-cAMP. (**C**) Western blot and densitometric analysis of the fibrosis-associated genes Col1A1 and αSMA in lung (top) and skin (bottom) SSc MFs following treatment with forskolin (20 µM) or db-cAMP (1 mM) for 96 h. (**D**) αSMA stress fibers were identified by immunofluorescence microscopy using an anti-αSMA-FITC-conjugated antibody (utilizing the same protocol in **C**). Nuclei were stained with DAPI. Data points represent distinct patient-derived cell lines. Significance for densitometric data in **C** (*n* = 6) was determined by one-way ANOVA. \**P* < 0.05, \*\**P <* 0.01, \*\*\**P* < 0.001 and \*\*\*\**P <* 0.0001.

### PGE_2_ promotes dedifferentiation in SSc lung, but not skin, MFs

Using publicly available single-cell RNA sequencing databases from SSc patients (24, 25), we examined the abundance of all canonical Gαs-coupled (cAMP-stimulatory) GPCR transcripts expressed in both lung and skin fibroblasts (Figure 2A, Supplemental Figure 2A). For this analysis, fibroblasts were identified by expression of *COL1A1, PDGFRA* and *FN1*. Relative abundance of these Gαs GPCRs was similar in both lung and skin cells. Transcripts encoding the two Gαs-coupled PGE_2_ receptors *PTGER2* (EP2) and *PTGER4* (EP4) exhibited the highest combined abundance in both lung and skin cells, followed by those encoding the prostacyclin receptor *PTGIR* (IP) and then the parathyroid hormone *PTHR1*. Previous work by our laboratory and others has demonstrated that activation of EP2/EP4 via their endogenous ligand PGE_2_ induces dedifferentiation of MFs derived from either normal or IPF lung (13, 17, 26). We therefore sought to compare the actions of PGE_2_ in SSc patient-derived lung and skin MFs, hypothesizing that its cAMP potentiating effects would likewise promote their dedifferentiation. To assess the effect of PGE_2_ in these cells, we used the same protocol detailed in Figure 1A, employing a supraphysiologic dose of PGE_2_ (1.0 μM) to maximize its effects. As expected, treatment of SSc lung MFs resulted in a significant reduction in both αSMA and collagen protein (Figure 2B) and transcript (Supplemental Figure 2C). Moreover, PGE_2_ promoted the complete disassembly of organized αSMA stress fibers (Figure 2C) and induced similar morphologic changes to SSc lung MFs dedifferentiated with forskolin and db-cAMP (Figure 1C, top panel). Unexpectedly, SSc skin MFs did not show a significant reduction in either αSMA or collagen and some patient-derived lines actually exhibited increases in these fibrosis-associated proteins (Figure 2B, Supplemental Figure 2B). Accordingly, the organized αSMA stress fibers and broad stellate morphology of these PGE_2_-resistant skin MFs were retained (Figure 2C, bottom panel).

**Figure 2.**
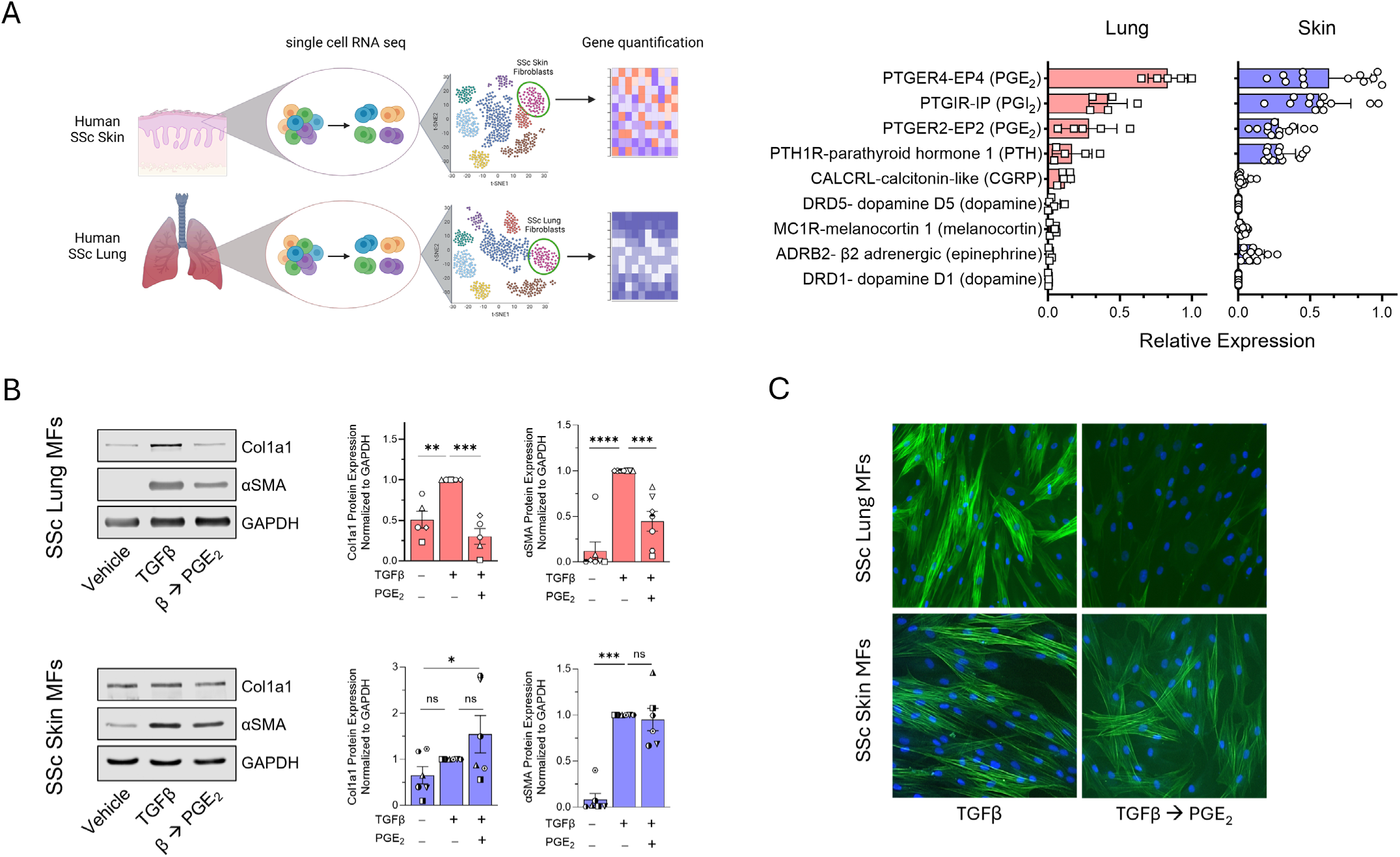
PGE_2_ promotes dedifferentiation in lung, but not skin, MFs. (**A**) Illustration (left) depicting the cellular source and method of acquisition for single-cell RNA sequencing data utilized from publicly available databases followed by subsequent gene expression analysis to identify fibroblast populations. Bar charts (right) displaying the baseline relative gene expression of the most highly expressed Gαs GPCR genes in SSc lung and skin fibroblasts from public single-cell sequencing databases. (**B**) Western blot analysis and densitometric analysis for the fibrosis-associated genes Col1A1 and αSMA from SSc lung and skin MF lysates following their treatment with PGE_2_ (1 µM) for 96 h. (**C**) αSMA stress fibers were identified by immunofluorescence microscopy using an anti-αSMA-FITC-conjugated antibody (using the same protocol in **C**). Nuclei were stained with DAPI. Data points represent distinct patient-derived cell lines. Significance for densitometric data in **B** (*n* = 5-7) was determined by one-way ANOVA. \**P* < 0.05, \*\**P <* 0.01, \*\*\**P* < 0.001 and \*\*\*\**P <* 0.0001.

### PGE_2_-mediated cAMP production in SSc lung MFs substantially exceeds that in SSc skin MFs

Despite the relatively high expression of Gαs-coupled EP2 and EP4 within both SSc lung and skin fibroblasts seen in the RNA sequencing databases, the discrepancy in the dedifferentiation capacity of PGE_2_ between MFs derived from the two organs stands out (Figure 2). This could indicate differences in EP receptor expression in TGFβ-differentiated MFs in culture (employed in our experiments) versus expression in fibroblasts determined by scRNA sequencing (reflected in the public databases). Alternatively, it could reflect increases in Gαi-coupled EP3 expression in MFs from skin compared to lung. Analysis of the same scRNA sequencing dataset in Figure 2A, showed that the expression of *PTGER3* (EP3) transcript was indeed the top expressed Gαi-coupled GPCR within SSc skin fibroblasts, with significantly higher expression than in SSc lung fibroblasts (Supplemental Figure 3A). We then measured the expression of EP2, EP3, and EP4 within the same patient lines used in Figures 1 and 2 to validate the scRNA sequencing data within MFs. Specifically, each fibroblast line was treated with TGFβ for 48 h to induce the MF phenotype and RNA was extracted for qPCR (Figure 3B). Among the Gαs-coupled receptor genes, EP2 expression was significantly higher in lung MFs while EP4 expression trended higher in skin. We also confirmed that the inhibitory Gαi-coupled EP3 transcript was expressed at markedly higher levels in skin than lung MFs (Figure 3B). To further justify our focus on EP2/EP4 agonism in response to PGE_2_, we measured *PTGIR* (IP) receptor expression within these same lines and found that its expression was dwarfed by that of EP2 and EP4 in both lung and skin (Supplemental Figure 3B). Additionally, TGFβ treatment of SSc fibroblasts from lung and skin had no significant effect on EP2, EP3, or EP4 transcript expression (Supplemental Figure 3C).

**Figure 3.**
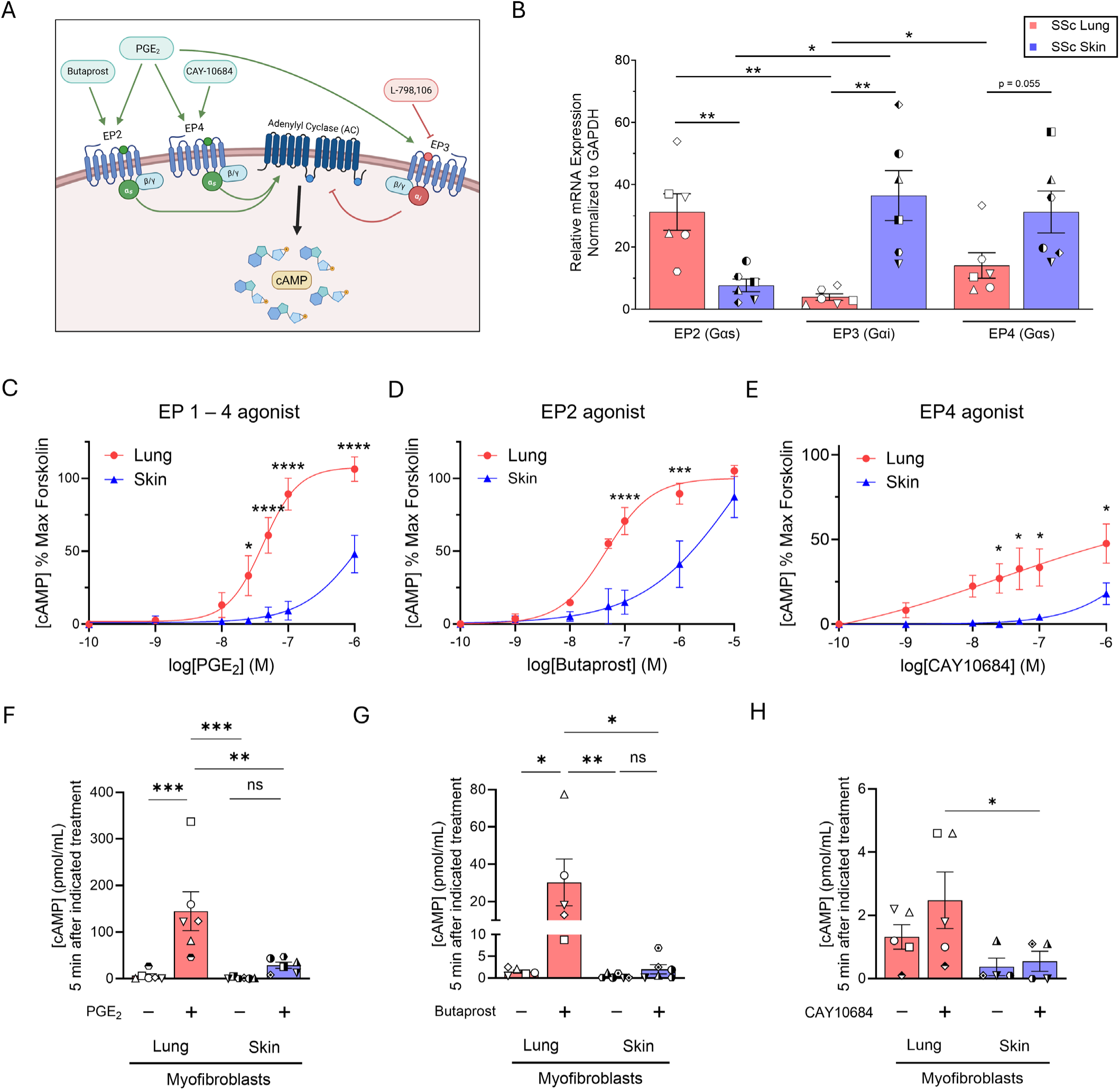
PGE_2_-mediated cAMP production in SSc lung MFs substantially exceeds that in SSc skin MFs. (**A**) Illustration depicting the AC activating effects of the Gαs-coupled GPCR agonists butaprost, PGE_2_ and CAY10684 at EP2 and/or EP4 resulting in an increase in cAMP levels. PGE_2_ exhibits inhibitory actions on AC by its ligation of Gαi-coupled EP3, while the EP3 antagonist L-798,106 prevents such AC inhibition. (**B**) qPCR data representing the relative gene expression of the *PTGER2*, *PTGER3* and *PTGER4* genes encoding EP2, EP3 and EP4, respectively, in SSc lung and skin MFs utilized in our experiments. Each symbol represents an individual patient-derived cell line. (**C-E**) Dose response curves of SSc lung and skin MFs treated for 2 h with PGE_2_ (designated EP1-4 agonist) (**C**), butaprost (**D**) or CAY 10684 (**E**), measured by cAMP-GLO assay; results are expressed relative to effects of forskolin as maximal stimulus. cAMP quantification in **C-E** is expressed as a proportion of maximum forskolin-mediated cAMP generation and was performed in the presence of the pan-PDE inhibitor IBMX (500 µM). (**F-H**) Intracellular cAMP concentration (pmol/mL) measured via ELISA in SSc lung and skin MFs following five-minute treatment with PGE_2_ (1 µM) (**F**), butaprost (1 µM) (**G**) or CAY10684 (1 µM) (**H**). Data points represent distinct patient-derived cell lines. Significance for comparisons in **B** was determined by 2-tailed unpaired or paired t-test where appropriate. Significance for cAMP data in **C-E** (*n* = 5) and **F-H** (*n* = 4-6) was determined by two-way ANOVA. \**P* < 0.05, \*\**P <* 0.01, \*\*\**P* < 0.001 and \*\*\*\**P <* 0.0001.

Since the dedifferentiating effects of PGE_2_ have been attributed to its ability to increase intracellular cAMP, mainly via EP2 (13), these transcript results suggest that the resistance of skin MFs to dedifferentiation could reflect their lower EP2 and/or higher EP3 expression compared to lung cells (13). To address these possibilities, we examined dose-dependent differences in cAMP production in SSc lung and skin MFs treated with PGE_2_, and also employed receptor-specific agonists/antagonists (Figure 3A schematic). cAMP levels were quantified in SSc MFs using the cAMP-GLO assay after treating cells with a cAMP modulating agent in the presence of the non-specific PDE inhibitor IBMX and PDE4 specific inhibitor Ro 20-1724 for 2 h. Inhibiting PDEs enables assessment of cAMP production alone without the confounding effects of its degradation. The cAMP values following treatment were then normalized to treatment over the same time span with forskolin to generate a dose-response curve for cells from each organ. SSc lung MFs exhibited significantly greater cAMP production in response to PGE_2_ treatment (referred to as “EP1-4 agonist”) than did skin MFs at doses greater than or equal to 25 nM (Figure 3C). The dose-response curve in skin cells (EC50 ∼1 µM) was shifted approximately 20-fold higher than in lung cells (EC50 ∼50 nM). While lung MFs were able to reach the same intracellular cAMP levels as forskolin at the 1 µM dose, skin MFs failed to reach even 50 percent of the maximum cAMP produced by forskolin. This result suggests that the impaired response to PGE_2_ in SSc skin MFs is likely related to an inadequate increase in intracellular cAMP. Moreover, the forskolin-mediated increase in intracellular cAMP was substantially greater in SSc lung MFs compared to skin as determined by ELISA (Supplemental Figure 3D), which directly measured cAMP concentration 5 min following AC stimulation in the absence of a PDE inhibitor. These results suggest that there is an even wider discrepancy between cAMP levels within the lung and skin dose curves in Figure 3C given that the values are normalized to the peak effects of forskolin. Differences in AC isoform transcript expression do not appear to account for the differences in cAMP levels following forskolin treatment in these two cell types (Supplemental Figure 3E).

We next tested the specific effects of selective EP2 and EP4 agonism on cAMP production. The selective EP2 agonist butaprost mirrored PGE_2_ in eliciting greater rises in cAMP in lung than skin MFs, with a similar difference in EC50 concentrations between the two. Only at a dose of 10 µM butaprost could the response in skin cells approach that of lung cells (Figure 3D). Finally, selective ligation of the EP4 receptor (via CAY10684) yielded a comparably diminished rise in cAMP in both cell types than did PGE_2_ or butaprost, reaching under 50% of the forskolin response in SSc lung MFs and 20% in skin (Figure 3E), suggesting that EP4 ligation results in substantially lower AC activity compared to EP2.

In a complementary approach, we measured precise cAMP concentration values for MFs via ELISA 5 min following treatment of MFs with 1 µM PGE_2_, butaprost, or CAY10684 – all in the absence of a PDE inhibitor. cAMP levels were significantly increased after treatment with PGE_2_ and butaprost in lung MFs, whereas skin MFs demonstrated a substantially blunted response to each of these treatments (Figure 3F and 3G). Notably, EP4 agonism yielded no significant increase in cAMP in either SSc lung or skin MFs (Figure 3H). Despite the non-significant increase in cAMP following treatment measured by ELISA in SSc skin MFs, butaprost (which did show cAMP accumulation over time by cAMP-GLO) elicited dedifferentiation in both lung and skin SSc MFs (Supplemental Figure 4A). The EP4 agonist CAY10684 resulted in reduced expression of αSMA and collagen within SSc lung MFs but failed to dedifferentiate SSc skin MFs (Supplemental Figure 4B).

Finally, we examined whether inhibition of the EP3 receptor could rescue PGE_2_-mediated cAMP production in SSc skin MFs. Following a 15-min pre-treatment with 250 nM of the EP3 antagonist L-978,106, MFs were treated with 1 µM PGE_2_. Surprisingly, EP3 inhibition did not increase cAMP production in either lung or skin MFs (Supplemental Figure 4C). Furthermore, pre-treatment with this EP3 inhibitor prior to PGE_2_ treatment failed to dedifferentiate SSc skin MFs or to potentiate the PGE_2_-elicited reduction in αSMA and collagen in SSc lung MFs (Supplemental Figure 4D). Taken together, these data indicate that PGE_2_ can robustly increase intracellular cAMP levels in lung, but not SSc skin MFs. Moreover, the reduced expression and/or reduced activity of EP2 within skin MFs – but curiously not the higher expression of EP3 – likely contributes to their reduced ability to generate cAMP as compared to lung MFs. It is also notable that EP4 appears to have a relatively minor role in cAMP production within both SSc lung and skin MFs relative to EP2.

### PDE4 inhibition augments PGE_2_-mediated cAMP production and promotes dedifferentiation in SSc lung and skin MFs

Although our data suggest that the distinct effects of PGE_2_ on SSc lung and skin MFs are partially explained by differences in E prostanoid receptor expression/activity, differences in cAMP degradation may also play a role. Endogenous PDEs function as a brake on cyclic nucleotide signaling by hydrolyzing the phosphodiester bond and converting the nucleotide monophosphate into a non-cyclized form (Figure 4A schematic). While there are eight PDE families that hydrolyze cAMP, fibroblasts are known to express high levels of PDE4 (27–31). The aforementioned single-cell databases revealed that PDE4 is indeed among the highest expressed PDE family within SSc lung and skin fibroblasts (Supplemental Figure 5A). PDE4 inhibition has been shown to dedifferentiate normal human skin fibroblasts (32) and prevent the differentiation of lung fibroblasts (28, 33–35), but its dedifferentiation capacity has not been assessed and compared within diseased lung and skin fibroblasts. We therefore hypothesized that pre-treatment with a PDE4 inhibitor would augment the effects of PGE_2_, thereby enabling it to dedifferentiate SSc skin MFs. We also hypothesized that treatment with a PDE4 inhibitor alone would augment any endogenous cAMP production from autocrine/paracrine sources and thereby be sufficient by itself to promote dedifferentiation of SSc MFs in culture.

**Figure 4.**
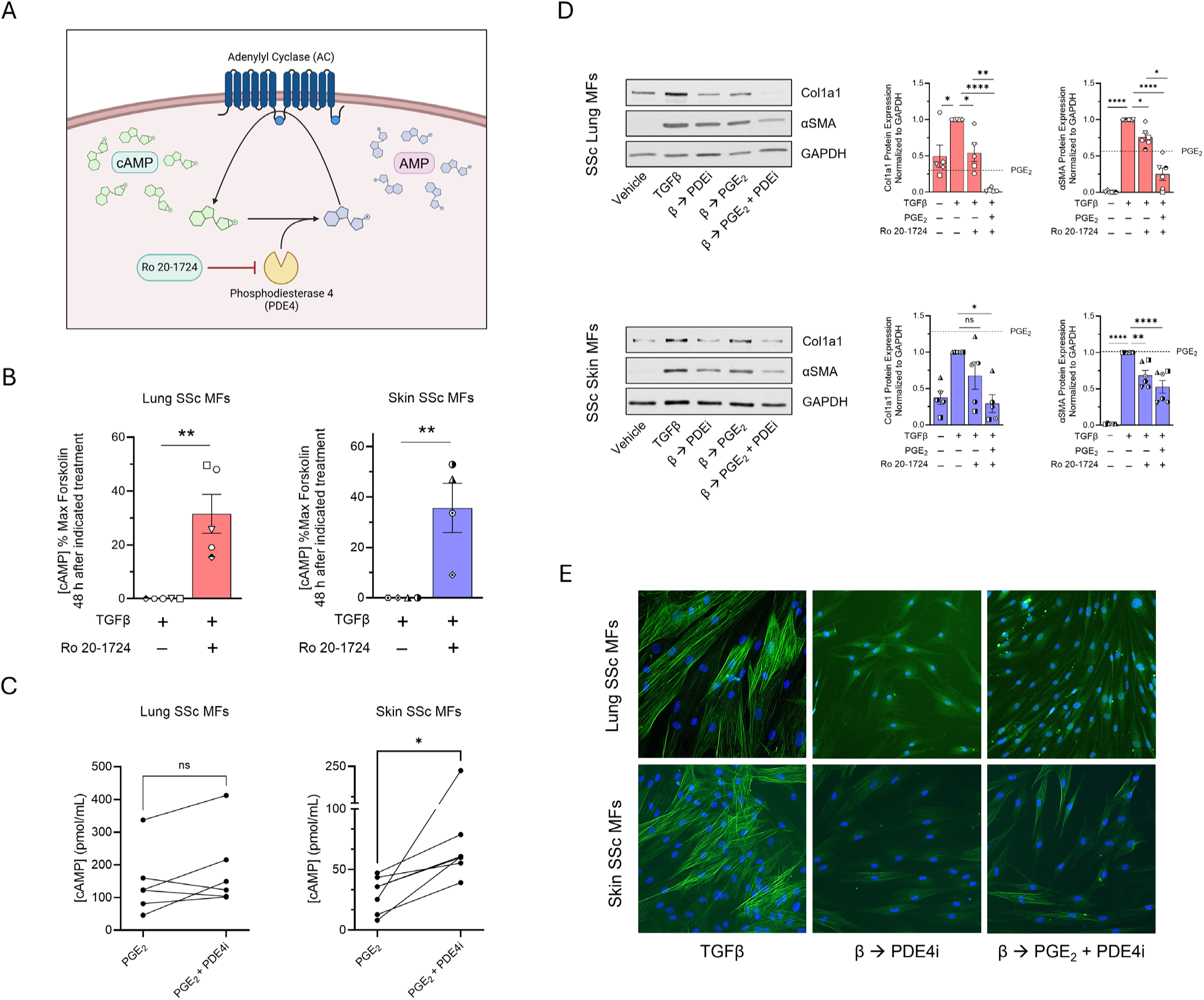
PDE4 inhibition augments PGE_2_-mediated cAMP production and promotes dedifferentiation in SSc lung and skin MFs. (**A**) Schematic depicting the action of PDE4 degradation of cAMP and its inhibition by the PDE4 inhibitor Ro 20-1724. (**B**) Intracellular cAMP production, measured by cAMP GLO, in SSc lung and skin MFs with and without 48 h incubation with Ro 20-1724 (100 µM); results are expressed relative to effects of forskolin as maximal stimulus. (**C**) Intracellular cAMP production, measured by ELISA, in SSc lung and skin MFs following PGE_2_ (1 µM) treatment for 5 min with and without 15 min pre-treatment with Ro-1724 (100 µM). (**D**) Western blot and densitometric analysis of the fibrosis-associated genes Col1A1 and αSMA following SSc lung and skin MF treatment with PGE_2_ (1 µM) for 96 h, with or without a 15 min pre-treatment of Ro-1724 (100 µM). Line depicts previous PGE_2_ treatment in Fig. 2B. (**E**) αSMA stress fibers were identified by immunofluorescence microscopy using an anti-αSMA-FITC-conjugated antibody (using the same protocol in **D**. Nuclei were stained with DAPI. Significance for data in **B-C** (*n* = 4-7) was determined by 1-tailed paired t-test. Significance for data **D-E** (*n* = 5) was determined by one-way ANOVA. \**P* < 0.05, \*\**P <* 0.01, and \*\*\*\**P <* 0.0001.

To assess each hypothesis, we first measured cAMP levels by ELISA following treatment with the global PDE4 inhibitor Ro 20-1724 in the presence or absence of PGE_2_. PDE4 inhibition alone had no significant effect on cAMP levels at 5 min as measured by ELISA (Supplemental Figure 5B), reflecting the lack of any GPCR-mediated activation of AC within this brief time window. However, when measured after 48 h via cAMP-GLO assay (without a non-specific PDE inhibitor), PDE4 inhibition alone produced substantial increases in cAMP (Figure 4B). This implied that lung and skin MFs indeed generated and secreted endogenous mediators at a basal tone capable of ligating Gαs-coupled GPCRs and activating AC in culture, and provided the rationale to test the dedifferentiating capacity of Ro 20-1724 by itself. Additionally, PDE4 inhibition potentiated the cAMP production in response to PGE_2_ at 5 min in SSc skin MFs (Figure 4C), suggesting that PDE4-mediated cAMP degradation may also contribute to PGE_2_ resistance within SSc skin MFs.

We evaluated the dedifferentiating effects of PDE4 inhibition on MFs, as depicted in the protocol in Figure 1A. PDE4 inhibition alone reduced αSMA protein and collagen in lung MFs over 96 h while only significantly reducing αSMA protein in skin (Figure 4D). However, when combined with PGE_2_, PDE4 inhibition significantly reduced collagen protein at 96 h compared to PGE_2_-alone in skin MFs. Moreover, PDE4 inhibition alone and in combination with PGE_2_ stimulated the breakdown of αSMA stress fibers and induced morphologic changes consistent with MF dedifferentiation in both organs (Figure 4E).

The ability of PDE4 inhibition to increase intracellular cAMP and elicit dedifferentiation in both SSc lung and skin MFs confirms that both cell types constitutively generate endogenous Gαs-coupled GPCR ligands and that baseline PDE4 tone constrains cAMP signaling in both. Furthermore, the ability of PDE4 inhibition to augment PGE_2_-mediated cAMP production to a greater extent in skin than lung SSc MFs and promote their dedifferentiation strongly suggests that PDEs (particularly PDE4 enzymes) exert substantial influence on cAMP signaling within these cells.

### cAMP-mediated p38 inhibition promotes dedifferentiation of lung and skin MFs

cAMP is known to interact with each of the three MAPK protein families: p38, ERK, and JNK. The profibrotic actions of TGFβ likewise influence MAPK activity via phosphorylation (36). Previous work by our laboratory has demonstrated that the profibrotic effects of TGFβ depend upon its activation of p38 via phosphorylation at Thr180/Tyr182 (37, 38). Additionally, cAMP-mediated dephosphorylation of p38 was shown to promote mouse and human lung MF dedifferentiation (7). We thus sought to verify whether cAMP signaling promotes inhibition of p38, ERK, and/or JNK within SSc lung and skin MFs. To assess the effect of cAMP on MAPK function, we measured phosphorylated p38, ERK, and JNK levels in SSc MFs treated with forskolin. In both lung and skin MFs, forskolin treatment significantly reduced p-p38 expression while having no effect on ERK (Figure 5A and 5B). Interestingly, p-JNK was significantly reduced in SSc skin MFs but not in lung. As p38 dephosphorylation was a common effect of forskolin treatment and it has been shown separately to influence lung and skin fibroblast phenotypes (7, 36), we hypothesized that p38 inhibition was responsible for the dedifferentiating effects of cAMP in both SSc lung and skin MFs. To test its capacity to do so, lung and skin MFs were treated with the pharmacologic p38 inhibitor SB203580 for 48 and 96 h to measure Col1A1 and αSMA transcript and protein, respectively. SB203580 was able to significantly reduce these MF markers in both lung and skin (Figure 5C, Supplemental Figure 6). Furthermore, immunofluorescence microscopy revealed that SB203580 promoted the complete dissolution of organized αSMA stress fibers (Figure 5D) and cell shape changes similar to those following forskolin treatment (Figure 1). We conclude that p38 is the downstream MAPK most likely to account for the dedifferentiating effects of cAMP in both SSc lung and skin MFs.

**Figure 5.**
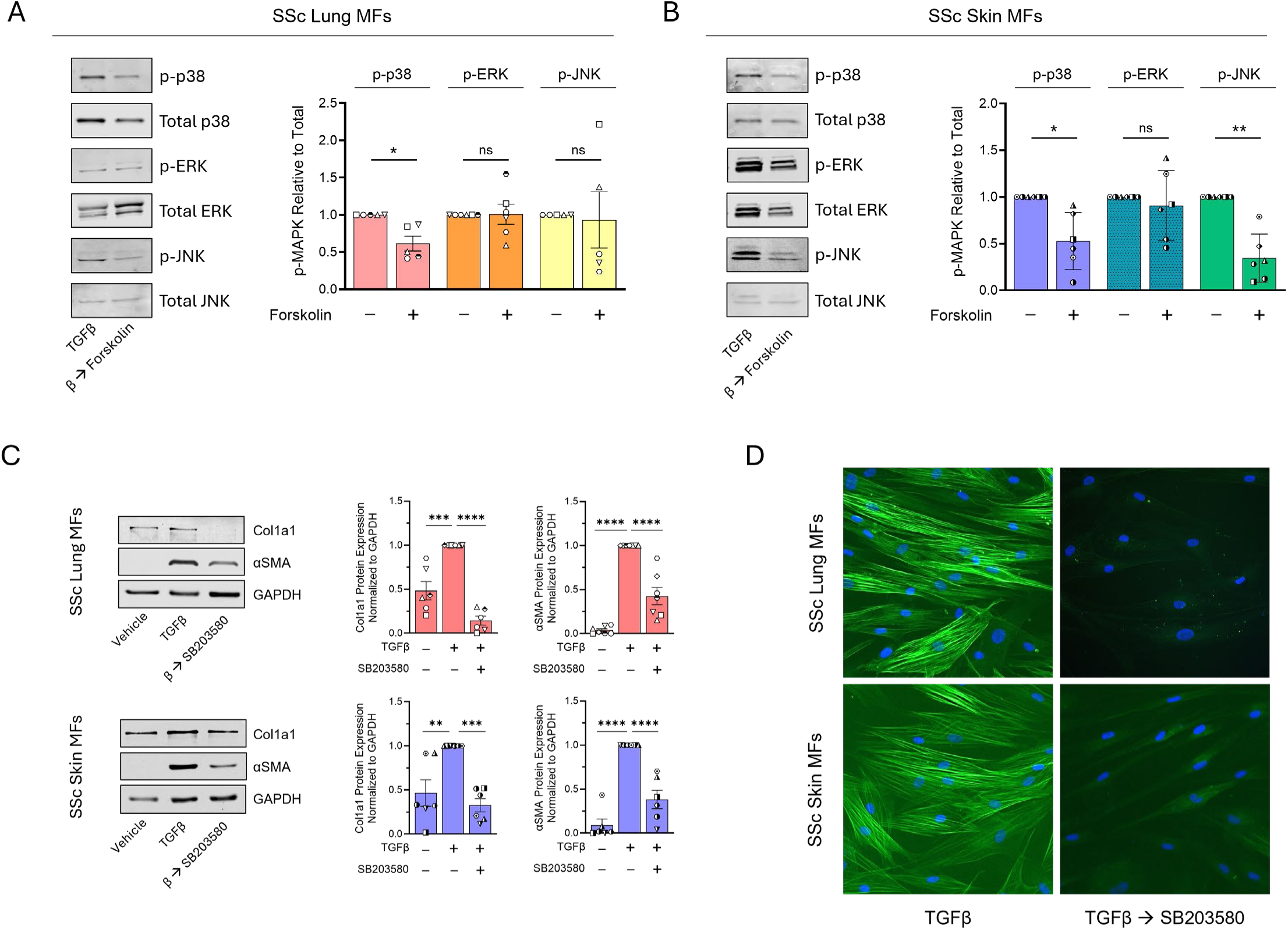
cAMP-mediated p38 inhibition promotes dedifferentiation of lung and skin MFs. (**A, B**) Western blot and densitometric analysis of the phosphorylated proteins p-p38, p-ERK and p-JNK following SSc lung (**A**) and skin (**B**) MF treatment with forskolin (20 µM) for 6 h. Phosphorylated proteins were normalized to total level of their respective proteins. (**C**) Western blot and densitometric analysis of the fibrosis-associated genes Col1A1 and αSMA following SSc lung and skin MF treatment with SB203580 (20 µM) for 96 h. (**D**) αSMA stress fibers were identified by immunofluorescence microscopy using an anti-αSMA-FITC-conjugated antibody (using the same protocol in **C**). Nuclei were stained with DAPI. Data points represent distinct patient-derived cell lines. Significance for densitometric data (*n* = 5-7) in **A** and **B** was determined and by 2-tailed paired t-test and by one-way ANOVA in **C**. \**P* < 0.05, \*\**P <* 0.01, \*\*\**P* < 0.001 and \*\*\*\**P <* 0.0001.

### p38α inhibition promotes SSc lung and skin MF dedifferentiation

The ultimate influence of each of the four p38 isoforms – p38α, p38β, p38γ and p38δ – depends upon their tissue restriction, spatiotemporal location within cells, and cellular context (39). We were therefore interested in determining the dominant isoform whose inhibition by cAMP promotes SSc lung and skin MF dedifferentiation. To assess the relative transcript abundance of each p38 isoform within SSc lung and skin MFs, we performed qPCR and found that in MFs from both organs p38α was the highest expressed isoform followed by p38γ, with minimal expression of p38β and δ (Figure 6A). We then treated MFs with the p38α-specific inhibitor VX-702 to assess the degree to which targeted inhibition of this isoform could recapitulate the dedifferentiating effects of the pan-p38 inhibitor SB203580. We again measured Col1A1 and αSMA transcript and protein 48 and 96 h after treatment, respectively. Similar to SB203580, VX-702 significantly reduced the expression of αSMA and collagen in SSc MFs from both organs (Figure 6B, Supplemental Figure 7) with a comparable reduction in αSMA stress fibers (Figure 6C). Notably, these results parallel our previous results demonstrating that cAMP-mediated dedifferentiation of human and mouse lung MFs was dependent upon inhibition of p38α (7). These findings therefore strongly suggest that p38α is the dominant profibrotic p38 isoform whose inhibition by cAMP likely accounts for the dedifferentiation of SSc lung and skin MFs.

**Figure 6.**
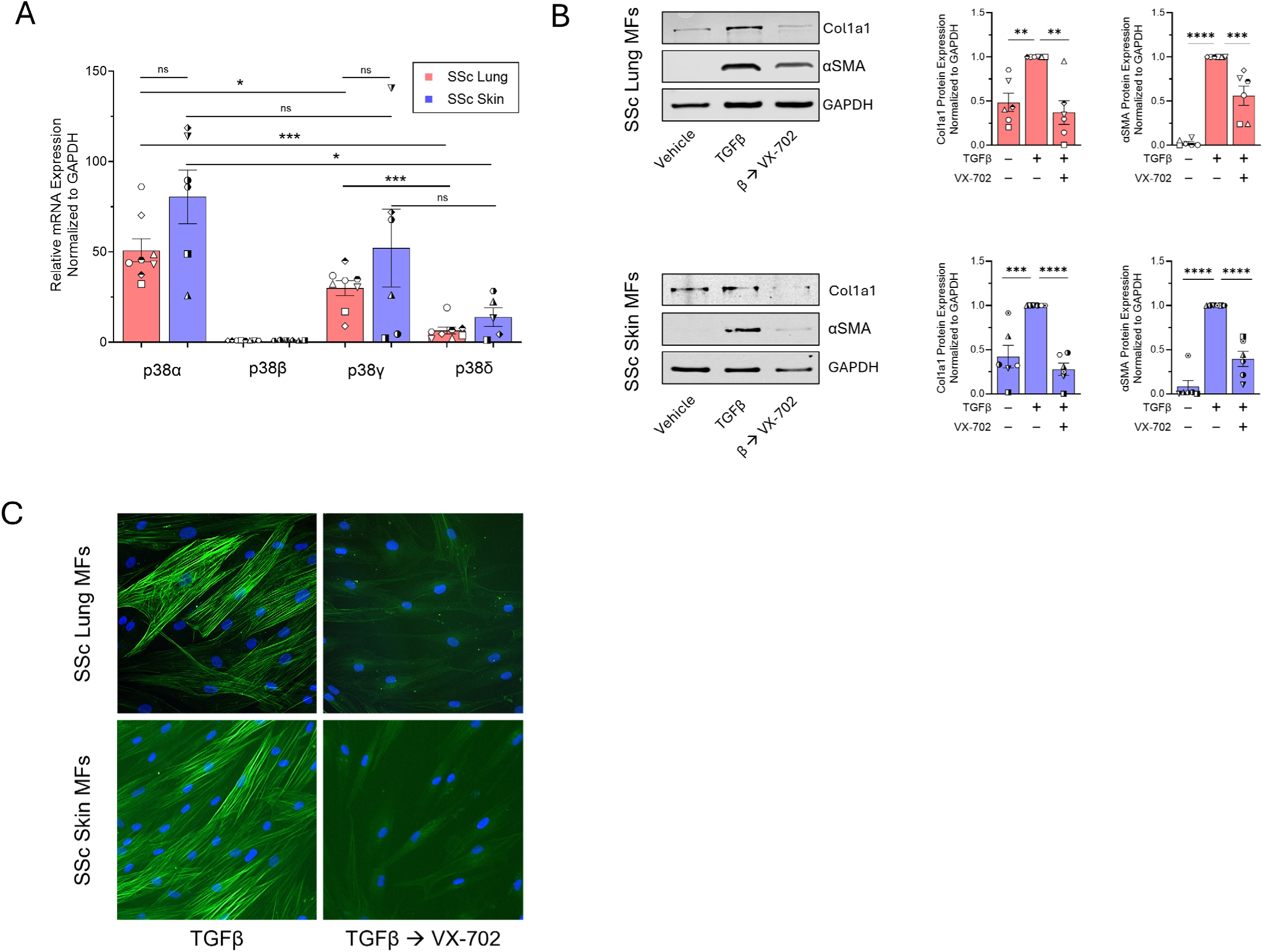
p38α inhibition promotes SSc lung and skin MF dedifferentiation. (**A**) qPCR data representing the relative expression of the genes encoding p38α, p38β, p38γ and p38δ in SSc lung and skin MFs. (**B**) Western blot and densitometric analysis of the fibrosis-associated genes Col1A1 and αSMA following SSc lung and skin MF treatment with the isoform-specific p38α inhibitor VX-702 (50µM) for 96 h. (**C**) αSMA stress fibers were identified by immunofluorescence microscopy using an anti-αSMA-FITC-conjugated antibody (using the same protocol in **B**). Nuclei were stained with DAPI. Data points represent distinct patient-derived cell lines. Significance for data in **A** (*n* = 8) was determined by 2-tailed unpaired or paired t-test where appropriate and by one-way ANOVA in **B** (n = 6). \**P* < 0.05, \*\**P <* 0.01, \*\*\**P* < 0.001 and \*\*\*\**P <* 0.0001.

## DISCUSSION

There are currently no approved drug treatments that reverse organ fibrosis in diseases such as SSc. It is now well recognized that deactivation and clearance of collagen-producing MFs are required for spontaneous fibrosis resolution in animal models (9, 10, 40), and these are therefore desirable features of effective antifibrotic therapies in human disease. Indeed, molecular mediators that induce the phenotypic dedifferentiation of MFs *in vitro* have been linked to their deactivation and clearance *in vivo* (7, 8, 11). In particular, the second messenger cAMP has been demonstrated to robustly disable numerous fibroblast functions (14, 41, 42) and to promote MF dedifferentiation (13, 17, 43). Intracellular potentiation of cAMP has recently garnered renewed interest as an antifibrotic therapy in IPF following the promising phase 2 results of the PDE inhibitor nerandomilast (44). Here we demonstrated that increases in intracellular cAMP within lung and skin MFs from patients with SSc induced their dedifferentiation through inhibition of the MAPK p38α. Though SSc MFs from the two organs exhibited similar phenotypic transitions in response to increased cAMP or direct inhibition of p38α, SSc skin MFs displayed resistance to the endogenous cAMP-stimulating mediator PGE_2_ – failing to substantially increase cAMP levels or dedifferentiate. Importantly, resistance to PGE_2_ in skin MFs was abolished by co-treatment with a PDE4 inhibitor and dedifferentiation of both lung and skin MFs was accomplished following treatment with this PDE4 inhibitor alone.

cAMP is a tightly regulated and highly conserved regulator of diverse cellular functions utilized by all living organisms (45). Its intracellular biosynthesis is almost exclusively accomplished by membrane-bound ACs whose activity is modulated through ligation of stimulatory or inhibitory GPCRs (46). Assessment of single-cell databases revealed expression of over eighty transcripts encoding distinct cAMP-modulating GPCRs within human SSc lung and skin fibroblasts (25, 47). Such a surfeit of receptors, the vast majority of which are activated by distinct ligands, ostensibly permits these cells to tailor the timing and magnitude of cAMP production to physiologic context and cellular requirements. Conversely, the intracellular half-life of cAMP is dependent upon the activity of several PDE enzyme families and gene isoforms (48) – whose spatial distribution can further influence and localize cAMP signaling within subcellular domains (49). Ultimately, upon reaching sufficient levels, cAMP exerts its effects by activating protein kinase A (PKA), the exchange protein activated by cAMP (Epac), and/or cyclic nucleotide-dependent ion channels (14, 50, 51). Although this work and that of several prior studies have shown that cAMP exhibits robust antifibrotic effects within tissue MFs (52–56), the complex cell- and tissue-specific nature of cAMP rheostasis must be acknowledged and considered when designing and implementing therapeutic strategies for fibrotic disease. This is especially true for SSc, in which multiple organs exhibit fibrosis simultaneously.

The distinct responses of SSc lung and skin MFs to PGE_2_ exemplifies the complexities of cAMP signaling within fibroblasts across tissues. The lesser capacity of skin than SSc lung MFs to generate cAMP in response to PGE_2_ (Figure 3C and 3F) is partially due to lower overall EP4 and especially EP2 activity (Figure 3D and 3E). It seemed plausible that the ability of the EP2 specific agonist butaprost (Supplemental Figure 3E), but not of PGE_2_ itself, to dedifferentiate skin MFs might be attributable to inhibition of AC via EP3 ligation by the latter. However, the inability of an EP3 inhibitor to potentiate either PGE_2_-mediated dedifferentiation or cAMP generation (Supplemental Figure 4C and 4D) argued against this notion. Thus, the ability of butaprost to induce dedifferentiation of MFs from both organs – in contrast to PGE_2_ – may reflect the additional confounding actions of the PGE_2_ receptors EP1 and EP4. EP4 has been shown to activate the Gαi subunit in certain cellular contexts (57, 58), thus inhibiting AC activity. Consistent with this possibility, the EP4 agonist CAY10684 did not significantly increase cAMP levels in skin MFs and failed to promote their dedifferentiation, yet accomplished both outcomes in lung (Figure 3E and Supplemental Figure 4B). Separately, EP1 is coupled with Gαq increasing cytoplasmic calcium levels – known to favor the MF phenotype (59, 60) – and can also activate Gαi (61). There is even evidence that EP2 can couple with Gαq under certain circumstances (62, 63), yet this putative profibrotic effect, if present, was dwarfed by the cAMP activating effects of EP2 in our study. It is also possible that the endogenous pharmacodynamic actions of PGE_2_ at the EP2 receptor may differ from those of the synthetic compound butaprost and influence cAMP kinetics in ways not captured by our assays. Finally, the magnitude of cAMP generation following direct activation of EP2 and/or AC may be additionally influenced by cell-specific factors that modulate the activity of each (46, 64). These results highlight the possible therapeutic advantage of individually agonizing a single receptor isoform (in this case, EP2) among the multiple isoforms that can be ligated by a molecule such as PGE_2_.

Distinct responses to forskolin and resistance to cAMP-activating ligands may reflect high baseline PDE activity in cells. Among the PDE families that exclusively target cAMP (PDE4, PDE7, PDE8), PDE4 was the highest expressed in both SSc lung and skin fibroblasts (Supplemental Figure 5A). We found that PDE4 inhibition increased PGE_2_-mediated cAMP generation and enabled it to elicit dedifferentiation in SSc skin MFs (Figure 4D). Moreover, PDE4 inhibition alone was sufficient to increase cAMP in both SSc lung and skin MFs (Figure 4B), suggesting that baseline autocrine/paracrine activity of fibroblasts from both organs stimulate Gαs-GPCRs in culture. However, the resulting cAMP levels induced by this cellular crosstalk may be below the threshold necessary to induce downstream signaling. One intriguing possible implication of this result for patients with SSc is the baseline presence of a suite of endogenous cAMP-generating ligands within the fibrotic microenvironments of the lungs and skin whose combined actions are nullified by continuous PDE4-mediated degradation of cAMP. Therapeutic inhibition of PDE4 would therefore be predicted to restore cAMP signaling in both lung and skin fibroblasts regardless of the underlying ligand(s) responsible for activating AC and generating cAMP in these cells. It is remarkable that inhibition of just one of the eight PDE enzyme families targeting cAMP could promote MF dedifferentiation. This is likely due in part to specific spatial distribution of PDE families within cells. The canonical cAMP target PKA is known to co-localize with PDE4 isoforms and post-transcriptionally activate them (65, 66), establishing a negative feedback loop. As the dedifferentiating effects of cAMP in MFs have been linked to the actions of PKA (13), inhibition of PDE4 may preferentially enhance PKA activity within signaling “microdomains” (67) and induce MF dedifferentiation. Such molecular events would therefore occur in spatial isolation from the effects of other active PDE enzyme families.

While a plethora of downstream signaling events have been attributed to the actions of cAMP (68), its influence on MAPK activity is of particular interest given the influence of this family of kinases on cellular phenotype and fate (69). The most consistent effect on the MAPKs following intracellular cAMP generation in our study was its dephosphorylation of p38 (Figure 5A and 5B). Pharmacologic inhibition of the p38α isoform within SSc lung and skin MFs was likewise sufficient to induce their dedifferentiation (Figure 6), mimicking the effects of intracellular cAMP. This finding supports our previous work linking the dedifferentiating actions of cAMP in lung MFs to induction of MAPK phosphatase 1 (MKP1) and its deactivation of p38α (7), suggesting a common antifibrotic role for this phosphatase within SSc lung and skin MFs. However, a previous study demonstrating increased inflammation and subsequent fibrosis within the skin following global MKP1 knockdown (70) emphasizes the cell-specific effects of downstream cAMP targets. Hence, therapeutic strategies that leverage cell-selective expression of GPCRs and/or specific PDE family isoforms may be necessary to achieve desired outcomes. With respect to the latter, studies to determine the organ- and cell-specific effects of the prospective new IPF therapy nerandomilast – a PDE4B-specific inhibitor (71) – will be necessary to determine its utility in SSc.

Taken together, our findings demonstrate the shared antifibrotic effects of cAMP-mediated p38α inhibition in human lung and skin fibroblasts from patients with SSc and also identify several important differences in the regulatory mechanisms that govern cAMP levels in these cells. We acknowledge specific limitations of this study including a lack of *in vivo* modeling and the inherent reductionist nature of studying fibroblasts in culture. Additionally, skin fibroblasts were obtained by biopsy upon informed consent from SSc patients exhibiting a spectrum of skin fibrosis (Supplemental Table 1), while lung fibroblasts were obtained exclusively from explanted lungs of patients with advanced stages of SSc-associated pulmonary fibrosis (Supplmental Table 2). Further validation of the pro-resolution effects of cAMP, investigation of different Gαs-coupled GPCR ligands and PDE4 inhibitors within multicellular systems such as organoids, precision cut tissue slices, and animal models of fibrosis, as well as patient-matched lung/skin samples, will be necessary to corroborate and strengthen our findings. Our results emphasize the therapeutic potential of cAMP signaling in lung and skin MFs and establish the importance of characterizing precisely how cAMP is regulated among fibroblasts from different tissues in SSc and in other fibrotic diseases.

## METHODS

### Reagents

Pharmacologic agents were reconstituted in DMSO to be stored as stock solutions at -80°C and were treated at working concentrations described in parentheses. Recombinant TGFβ purchased from R&D (7754-BH) was resuspended at a concentration of 2 ng/mL in filter-sterilized 1% BSA. The direct AC activator forskolin (20 µM) and the cAMP analog dibutyryl-cAMP (bucladesine sodium salt, 1 mM) were purchased from Selleckchem (S2449) and Cayman Chemicals (14408), respectively. PGE_2_ (1 µM); the EP2 agonist butaprost (1 µM); the EP4 agonist CAY10684 (1 µM); the EP3 antagonist L-798, 106 (250 nM); the PDE4 inhibitor Ro 20-1724 (100 µM); and the pan-PDE inhibitor IBMX (500 µM) were purchased from Cayman Chemicals (14010, 13740, 15966, 11129, 18272, 13347). The pan-p38 inhibitor SB203580 (20 µM) and p38α inhibitor VX-702 (50 µM) were purchased from Cayman Chemicals (13067) and Selleckchem (HY-10401), respectively.

### Cell culture

Lung and skin fibroblasts from separate groups of patients with SSc were obtained and employed as previously described (72–74). Deidentified demographic and clinical information for each patient whose skin or lung fibroblasts were utilized in this study are detailed in Supplemental Tables 1 and 2, respectively. Cells were cultured in a medium composed of low glucose DMEM (Invitrogen), 10% FBS (Hyclone), 100 units/mL penicillin, and 100 µg/mL streptomycin (both Invitrogen). Cell passages 4-8 were used for experiments. Upon reaching 70-80% confluence, cells were serum-starved in FBS-free DMEM overnight, followed by TGFβ treatment for 48 h to induce the MF phenotype. SSc lung and skin MFs were then utilized for experimental readouts as described. For each experiment, four to eight distinct lung or skin patient cell lines were utilized.

### qPCR

Cellular RNA was extracted via the RNeasy kit (Qiagen) to be analyzed for transcript expression. cDNA was synthesized from RNA extracts using the High-Capacity cDNA Reverse Transcription Kit (Applied Biosystems), then PCR products were amplified in Fast SYBR Green Master Mix (Invitrogen) and measured on the StepOne real-time PCR system (Applied Biosystems). Fold changes in transcript expression were normalized to the housekeeping gene GAPDH. All primer pair sequences used in this study are contained in Supplementary Table.

### Western Blot

Cells were lysed using RIPA buffer containing protease inhibitors (Roche Diagnostics, 11836153001) and a phosphatase inhibitor cocktail (EMD Biosciences, 524624 and 524625). Protein samples were separated by SDS-PAGE and then transferred onto either nitrocellulose or polyvinylidene difluoride (PVDF) membranes. The membranes were blocked with 5% BSA and subsequently incubated with primary antibodies, including mouse antibodies specific to αSMA (Dako), and GAPDH (Invitrogen) or rabbit antibodies against collagen 1a1 (Cell Signaling Technologies – CST, 91144), total p38 (CST, 9212), phospho-p38 (CST, 9211), total JNK (CST, 9252), phospho-JNK (CST, 9251), total ERK (CST, 9102), and phospho-ERK (CST, 4370). For experiments involving MAPKs (p38, ERK, JNK), the membranes were first probed for the phosphorylated form of the MAPK and then stripped and re-probed for the corresponding total MAPK, which served as a loading control. GAPDH was used as the loading control for all other Western blots. Proteins of interest and their respective loading controls were run on the same gel.

### cAMP Quantification by cAMP ELISA

SSc lung and skin fibroblasts were differentiated for 48 h with TFGβ as previously described. Cells were treated with forskolin, PGE_2_, butaprost, or CAY10684, then cAMP was assayed with the Cyclic AMP ELISA Kit (Cayman). Intracellular cAMP levels were measured with the Infinite M Plex (Tecan) and subsequently analyzed.

### cAMP quantification by cAMP-GLO

SSc lung and skin fibroblasts were plated on Biocoat Poly-D-Lysine 96-well plates (Corning) and differentiated with TGFB for 48 h as previously described. cAMP production was then analyzed following the cAMP-GLO Assay (Promega) protocol. Briefly, medium was removed and replaced with serum-free medium containing the PDE inhibitors Ro 20-1724 and IBMX. Then, cells were treated with forskolin (20 µM); various doses of PGE_2_, butaprost, or CAY10684; or left untreated for 2 h. The resulting luminescence was then measured on the Infinite M Plex (Tecan) and analyzed to determine their cAMP production compared to forskolin. While the cAMP-GLO assay allows intracellular cAMP to accumulate over 2 h in the presence of PDE inhibitors to obtain a maximal cAMP production estimate solely from GPCR stimulation, the cAMP ELISA measures cAMP generation after 5 min in the absence of any PDE inhibitors, allowing insights into a more biologically relevant assessment of cAMP biosynthesis. Primer pair sequences for each gene quantified by qPCR are displayed in Supplemental Table 3.

### Immunofluorescence microscopy

SSc lung and skin fibroblasts were seeded onto single-chamber slides and cultured, followed by serum starvation overnight. To induce fibroblast differentiation into MFs, TGFβ was added at a concentration of 2 ng/mL for 48 h. Cells were then treated with either a vehicle control or the specified agents to promote MF dedifferentiation. Afterward, the chamber slides were rinsed twice with cold PBS, fixed using freshly prepared 4% formaldehyde for 20 min, rinsed again with PBS, and quenched with 100 mM glycine for 15 min. Blocking and permeabilization were conducted by incubating the slides for 1 h in PBS containing 10% FBS and 0.1% Triton X-100 (Sigma-Aldrich). The staining process involved using the following primary antibodies: anti-αSMA-FITC (1:500; F3777, Sigma-Aldrich). For unconjugated primary antibodies, the fluorescent secondary antibody Cy3 (1:250; Jackson 711-166-152) was used for imaging. A mounting medium containing DAPI (Invitrogen) was employed to label nuclei. The slides were then imaged using a Revolve2 hybrid microscope (Echo).

### Single-cell RNA Sequencing Analysis

Raw single-cell RNA sequencing (scRNA-seq) data were obtained from the Gene Expression Omnibus (GEO) under accession numbers GSE212109 and GSE138669 for lung and skin, respectively. The lung dataset was subsetted to focus on five samples representing lung tissue from SSc-ILD patients (GSM6509488, GSM6509489, GSM6509490, GSM6509491, GSM6509497). 12 skin samples from SSc patients were selected from the skin dataset (GSM4115869, GSM4115871, GSM4115873, GSM4115877, GSM4115879, GSM4115881, GSM4115882, GSM4115883, GSM4115884, GSM4115887, GSM4115888, GSM4115889). Lung and skin sc-RNA sequenencing datasets were each handled separately in two different scripts. The raw feature-barcode matrices were loaded using the Read10X_h5 function in Seurat (v5.1.0). For each sample, a Seurat object was created using the CreateSeuratObject function, with a minimum threshold of 200 features and 3 cells per gene to filter out low-quality cells. The mitochondrial gene content was calculated using the PercentageFeatureSet function by identifying genes with the prefix “MT-” to estimate mitochondrial contamination. The individual Seurat objects were merged into a single object using the merge function. To ensure high-quality data, cells were further filtered based on several criteria: cells with fewer than 1,000 or more than 20,000 unique molecular identifiers (UMIs), cells expressing fewer than 1,000 genes, and cells with more than 20% mitochondrial content were excluded from downstream analysis. The data were then normalized using Seurat’s NormalizeData function, and highly variable features were identified with FindVariableFeatures. Data scaling was performed using ScaleData, followed by dimensionality reduction via principal component analysis (PCA) using RunPCA.

For integration of the individual samples composing the merged Seurat object, reciprocal PCA (RPCA) was used as an integration method through the IntegrateLayers function. The integrated data were used to construct a shared nearest neighbor graph (FindNeighbors), followed by clustering of cells at a resolution of 0.5 (FindClusters). UMAP (Uniform Manifold Approximation and Projection) was applied for visualization of clusters in two dimensions (RunUMAP).

To identify fibroblast populations, clusters expressing the marker genes *COL1A1*, *PDGFRA*, and *FN1* were visualized using FeaturePlot. Fibroblast populations were further subsetted based on their cluster identities (subset) and re-annotated by their original sample identity (Idents). Gene expression was aggregated across fibroblast populations using AggregateExpression, and the resulting counts per million (CPM) matrix was normalized using relative counts normalization (NormalizeData, normalization method = “RC”). Finally, the normalized CPM data were exported as a CSV file for further analysis.

### Statistics

Statistical analysis was performed using GraphPad Prism, version 10.1.1 (GraphPad Software). Experimental data are presented as the mean and were analyzed for significance by 1-way ANOVA with Tukey’s multiple-comparison test or paired/unpaired t-test, as indicated. A *P* value of <0.05 was considered significant. Unless otherwise noted, error bars represent standard error of the mean.

## AUTHOR CONTRIBUTIONS

JDB, SMF, and MPG designed the experiments. Experiments were performed by JDB and SMF. Data were analyzed by JDB and SMF. Patient derived SSc lung fibroblasts and SSc skin fibroblasts as well as intellectual contributions were provided by CFB and JV, respectively. The manuscript was written by JDB, SMF, and MPG. All authors reviewed the manuscript.

## ACKNOWLEDGEMENTS

We thank members of the Peters-Golden and Varga laboratories for their valuable input into this work and Swati Bhattacharyya for preparing and supplying the patient-derived SSc skin fibroblasts. This work was supported by a grant from the National Scleroderma Foundation (to SMF) and NIH grants K08 HL163178 (to SMF), K24 AR060297 (to CFB), R01 AR074997 (to JV), and R35 HL144979 (to MPG). All figure schematics were created with BioRender (biorender.com).

