## Supplemental Materials for "Distinct cAMP regulation in scleroderma lung and skin myofibroblasts governs their dedifferentiation via p38α inhibition"

Supplemental Figures and Tables

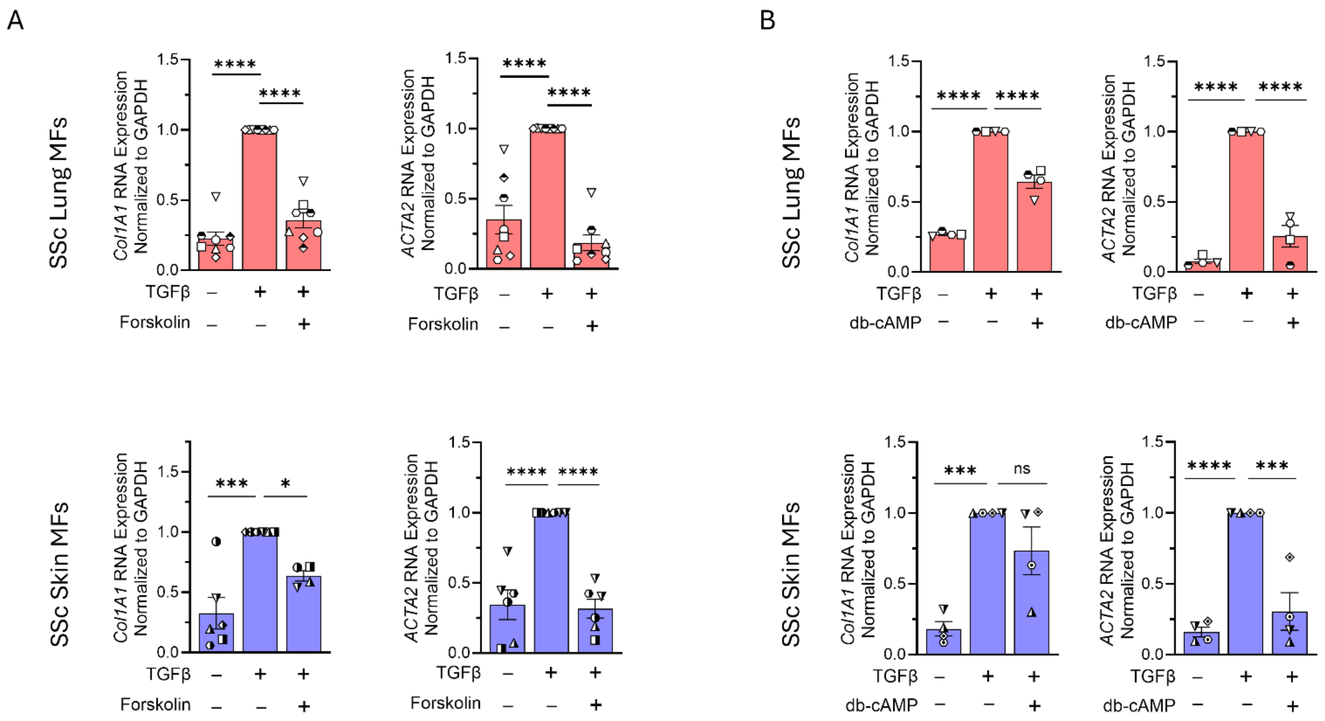


**Supplemental Figure 1.** (**A** - **B**) qPCR analysis of fibrosis-associated genes *Col1A1* and *ACTA2* in SSc lung and skin MFs following 48 h treatment with forskolin (20 µM) (**A**) or db-cAMP (1 mM) (**B**). Data points represent distinct patient-derived cell lines. Significance for qPCR data in **A** (*n* = 4-8) and **B** (*n* = 4) was determined by one-way ANOVA. **P* < 0.05, ****P* < 0.001 and *****P* < 0.0001.

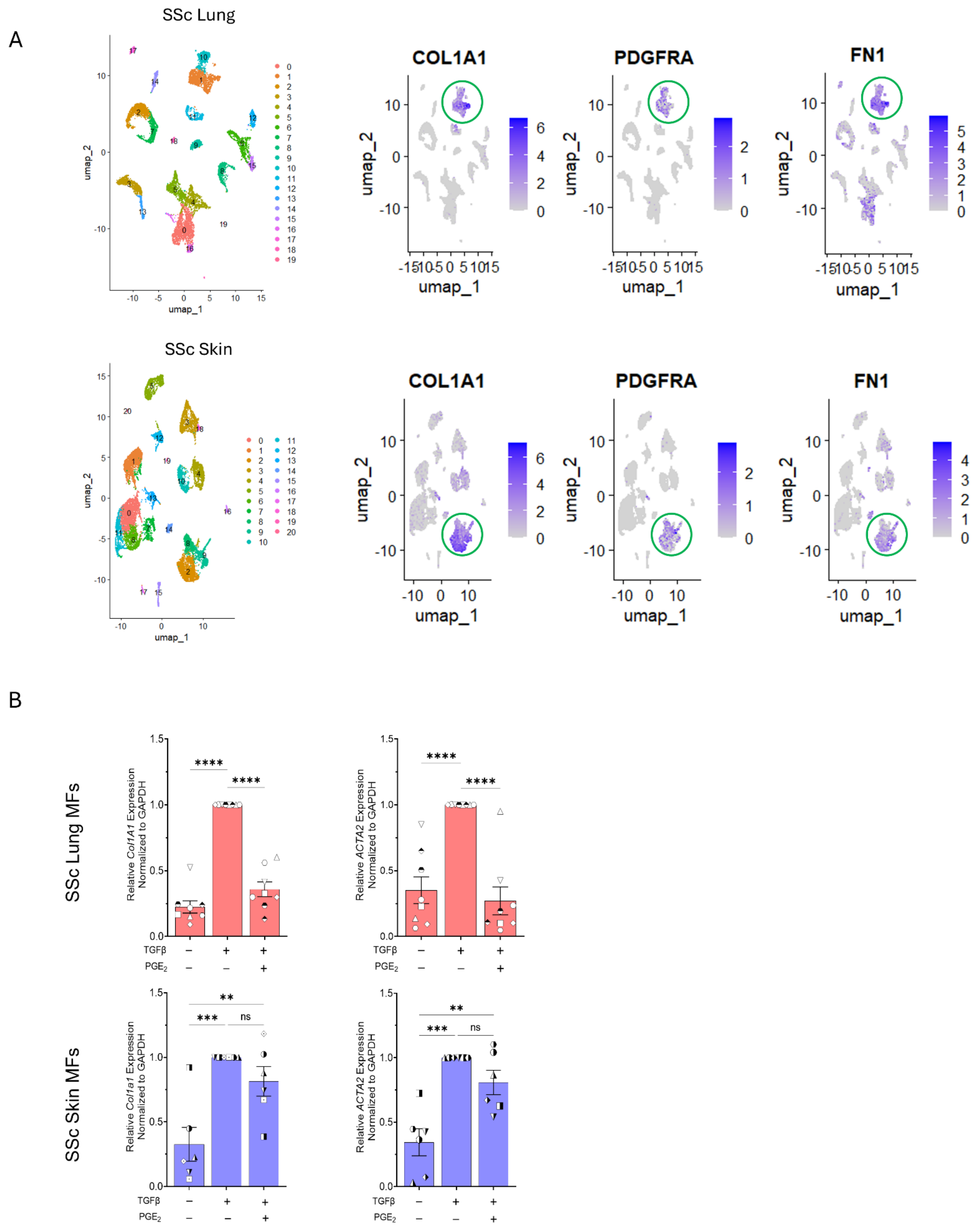

A
SSc Lung
umap_1
umap_2
COL1A1
PDGFRA
FN1
SSc Skin
B
SSc Lung MFs
SSc Skin MFs
Relative Col1A1 Expression Normalized to GAPDH
Relative ACTA2 Expression Normalized to GAPDH
Relative Col1a1 Expression Normalized to GAPDH
****
***
**
ns
TGFβ
PGE2

**Supplemental Figure 2.** (**A**) Illustration depicting the clustering of public single-cell RNA sequencing SSc skin and lung data. Fibroblast clusters were selected based on enrichment in profibrotic marker genes *Col1A1, PDGFRA*, and *FN1.* (**B**) qPCR analysis of fibrosis-associated genes *Col1A1* and *ACTA2* in SSc lung and skin MFs following 48 h treatment with $PGE_2$ (1 μM). Data points represent distinct patient-derived cell lines. Significance for qPCR data in **B** (n= 6-8) was determined by one-way ANOVA. *$P < 0.05$, **$P < 0.01$, ***$P < 0.001$ and ****$P < 0.0001$.

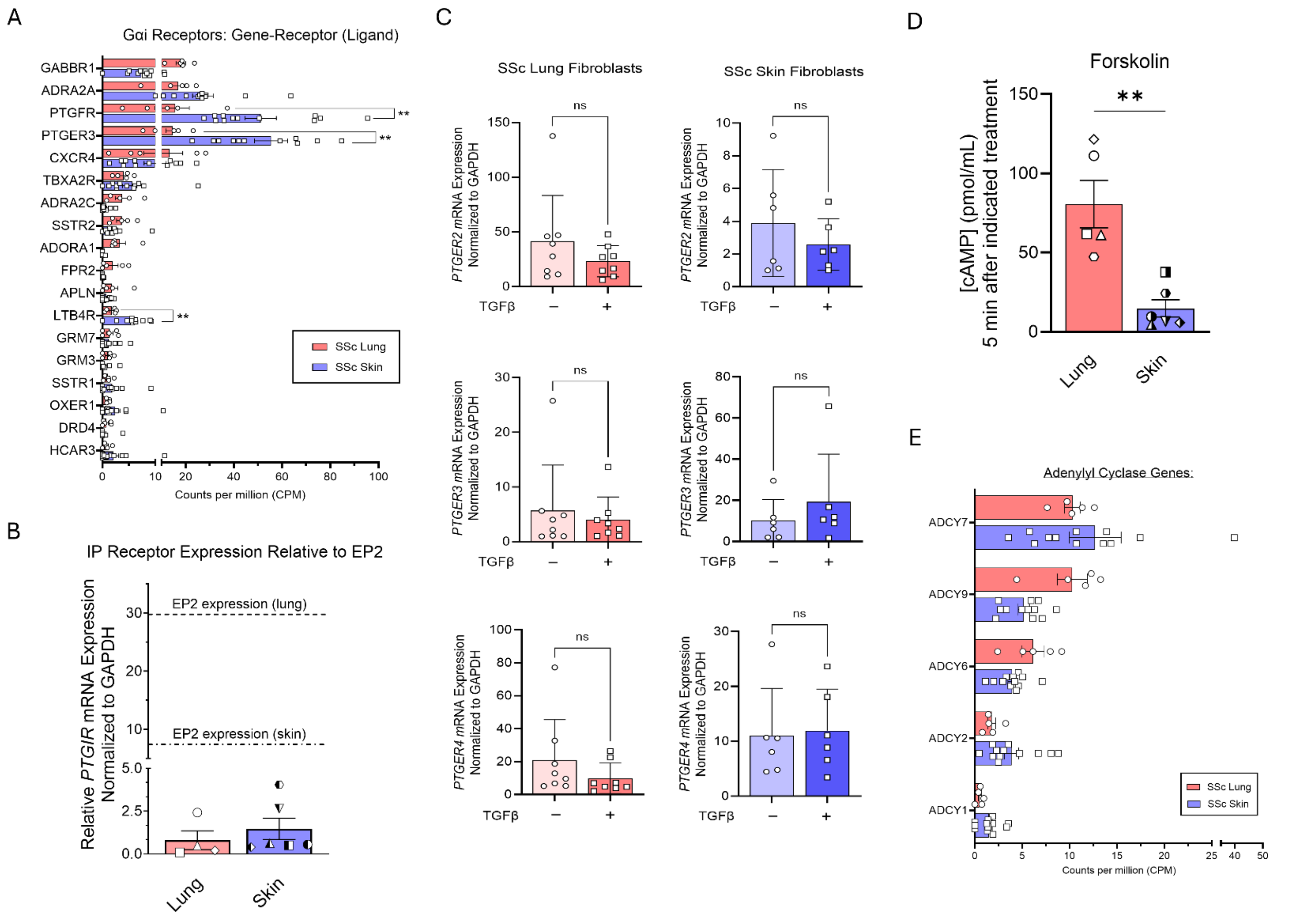


**Supplemental Figure 3.** (**A**) Bar chart illustrating the baseline gene expression in counts per million (CPM) of the most highly expressed Gαi GPCR genes in SSc lung and skin fibroblasts from public single-cell sequencing databases. (**B**) qPCR data representing the relative gene expression of the *PGTIR* gene encoding the IP receptor in SSc lung and skin MFs. Dashed line depicts relative EP2 gene expression level depicted in Figure 3B. (**C**) qPCR analysis of cAMP-modulating EP receptors in lung and skin SSc fibroblasts with or without TGFβ treatment for 48 h. (**D**) Intracellular cAMP concentration (pmol/mL) measured via ELISA in SSc lung and skin MFs following treatment with forskolin (20 µM) for 5 min. (**E**) Bar chart illustrating the baseline gene expression in counts per million (CPM) of the AC genes expressed in SSc lung and skin fibroblasts from public single-cell RNA sequencing databases. Data points represent distinct patient-derived cell lines. Significance for data in **A**, **C**, and **D** ($n$ = 5-12) was determined by 2-tailed unpaired t-test. *$P < 0.05$, **$P < 0.01$, ***$P < 0.001$ and ****$P < 0.0001$.

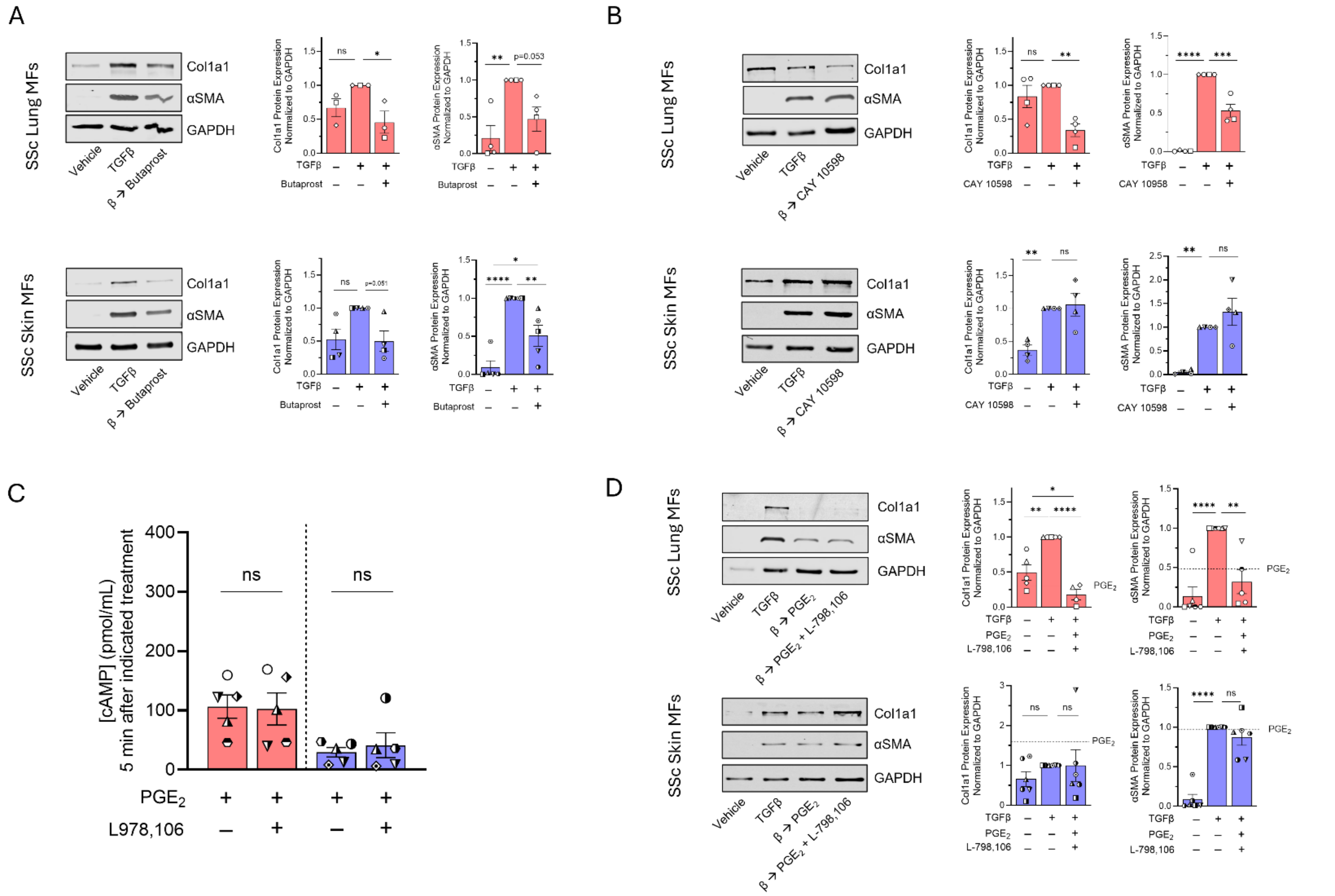


**Supplemental Figure 4.** (**A**) Western blot analysis of the fibrosis-associated genes Col1A1 and αSMA and densitometric analysis following SSc lung and skin MF treatment with butaprost (1 µM) for 96 h. (**B**) Western blot analysis of the fibrosis-associated genes Col1a1 and αSMA and densitometric analysis following SSc lung and skin MF treatment with CAY10598 (1 µM) for 96 h. (**C**) Intracellular cAMP concentration (pmol/mL) measured via ELISA in SSc lung and skin MFs following treatment with $PGE_2$ (1 µM) for 5 min, with or without a 15 min pre-treatment with the EP3 inhibitor L-798,106 (250 nM). (**D**) Western blot analysis of the fibrosis-associated genes Col1A1 and αSMA and densitometric analysis following SSc lung and skin MF treatment with $PGE_2$ (1 µM) for 96 h, with or without a 15 min pre-treatment with the EP3 inhibitor L-798,106 (250 nM). Dashed line depicts mean densitometry from previous $PGE_2$ treatment in Figure 2B. Data points represent distinct patient-derived cell lines. Significance for data in **C** was determined by 2-tailed paired t-test and by one-way ANOVA in **A,B, D** (*n* = 3-6). *$P < 0.05$, **$P < 0.01$, ***$P < 0.001$ and ****$P < 0.0001$.

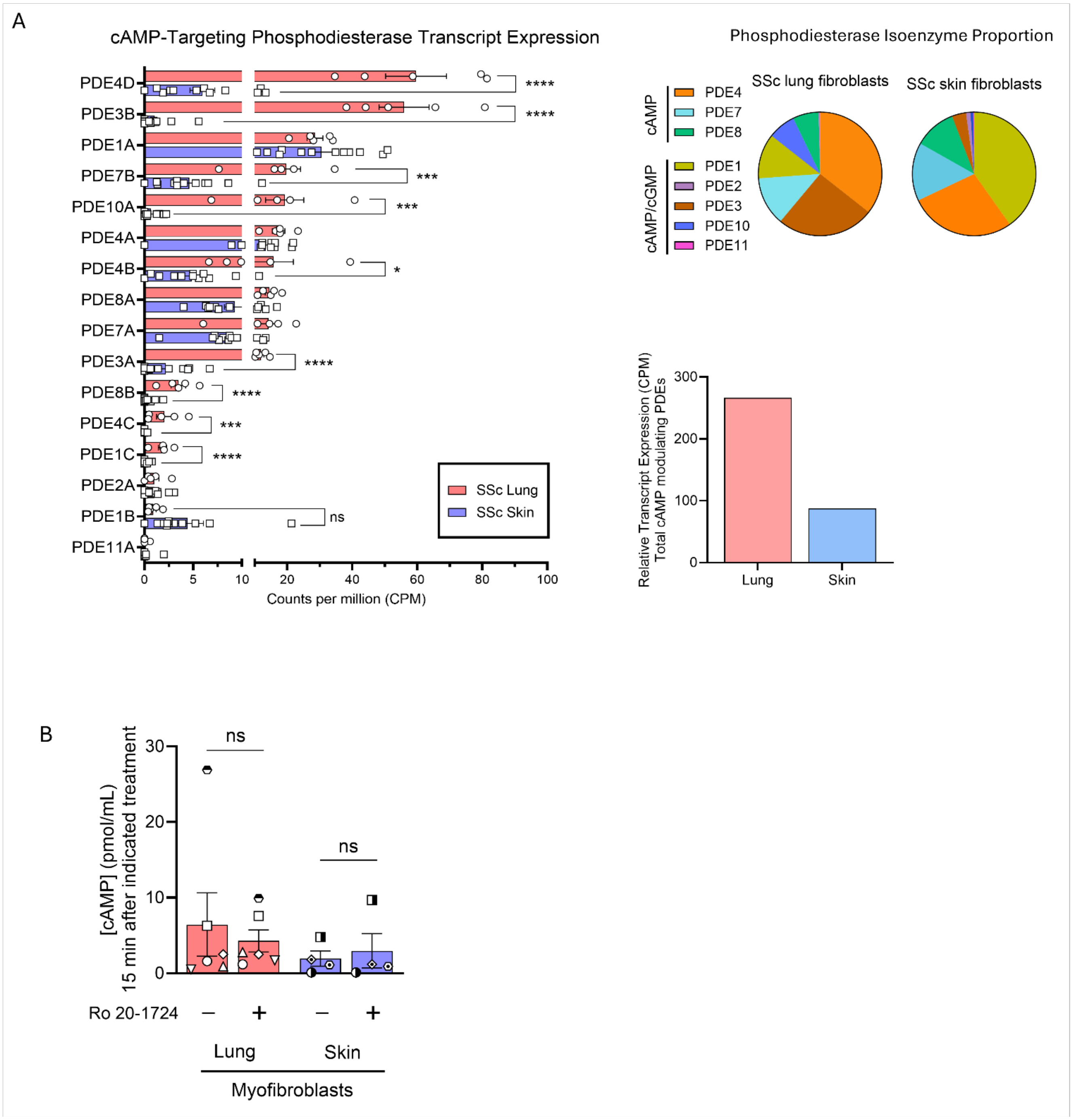


**Supplemental Figure 5.** (**A**) Bar chart illustrating the baseline gene expression in counts per million (CPM) of the PDE genes expressed in SSc lung and skin fibroblasts from public single-cell RNA sequencing databases (left). Pie chart representing the proportion of each PDE family compared to all PDE genes expressed in the fibroblast populations (top right). Bar chart demonstrating the total gene expression of cAMP modulating PDEs in CPM (bottom right). (**B**) Intracellular cAMP concentration in SSc lung and skin MFs before and after 15 min treatment with Ro 20-1724 (100 µM). Data points represent distinct patient-derived cell lines. Significance for B (n= 4-5) was determined by 1-tailed paired t-test.

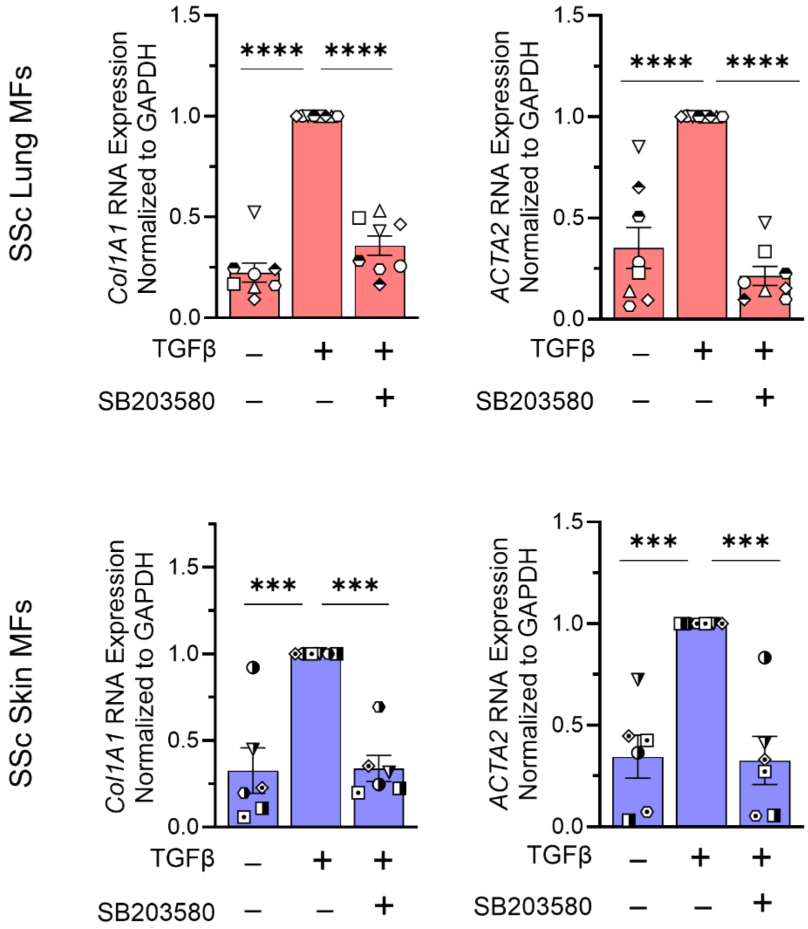


**Supplemental Figure 6.** qPCR analysis of fibrosis-associated genes *Col1A1* and *ACTA2* in SSc lung and skin MFs following treatment with SB203580 (20 µM) for 48 h. Data points represent distinct patient-derived cell lines. Significance for qPCR data (*n* = 6-8) was determined by one-way ANOVA. ***$P < 0.001$ and ****$P < 0.0001$.

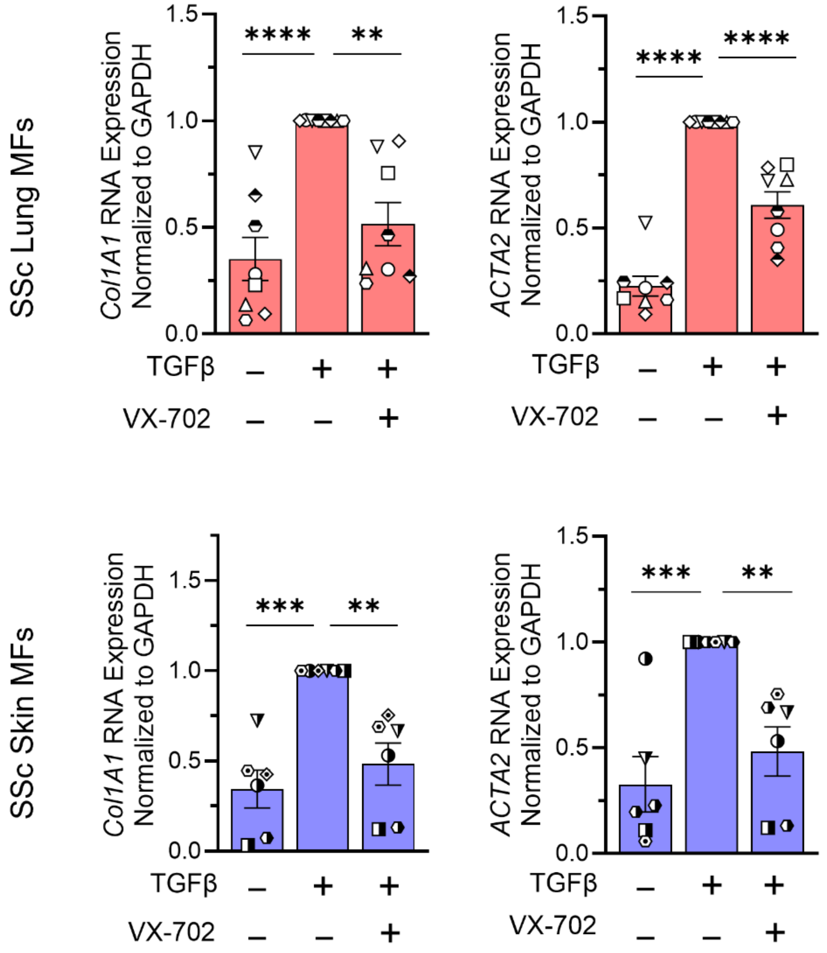


**Supplemental Figure 7.** qPCR analysis of fibrosis-associated genes *Col1A1* and *ACTA2* in SSc lung and skin MFs following treatment with VX-702 (50 μM) for 48 h. Data points represent distinct patient-derived cell lines. Significance for qPCR data (*n* = 6-8) was determined by one-way ANOVA. $^{**}P < 0.01$, $^{***}P < 0.001$ and $^{****}P < 0.0001$.

| Sample | Age | mRSS | Forearm mRSS | FVC (% pred) | TLC (% pred) | DLCO (% pred) |
|---|---|---|---|---|---|---|
| SPARC_SSc_21 | 49 | 8 | 1 | 79 | 93 | 60 |
| SPARC_SSc_22 | 51 | 9 | 1 | 90 | 109 | 84 |
| SPARC_SSc_23 | 37 | 47 | 2 | 63 | 99 | 25 |
| SPARC_SSc_25 | 45 | 12 | 2 | 100 | 98 | 101 |
| SPARC_SSc_29 | 50 | 33 | 2 | 70 | 83 | 72 |
| SPARC_SSc_33 | 49 | 13 | 2 | 49 | 62 | 50 |
| SPARC_SSc_34 | 64 | 28 | 2 | 48 | 52 | 41 |
| SSc_52 | 52 | - | - | - | - | - |

**Table 1** Patient demographics and disease-specific data. mRSS = modified Rodnan Skin Score. FVC = forced vital capacity, TLC = total lung capacity, DLCO = diffusion capacity. “-“ data not available.

| Sample | Age | Lung Pathology | FVC (% pred) | TLC (% pred) | DLCO (% pred) |
|---|---|---|---|---|---|
| SSc-24 | 45 | UIP/NSIP | 26 | - | 30 |
| SSc-25 | 51 | UIP/granuloma | 23 | 30 | - |
| SSc-26 | 57 | UIP | 33 | 33 | 26 |
| SSc-30 | 58 | UIP | 39 | 46 | 14 |
| SSc-59 | 42 | - | 18 | 30 | - |
| SSc-63 | 54 | - | 46 | 51 | 18 |
| SSc-74 | 49 | - | 56 | - | 24 |
| SSc-85 | 62 | - | 46 | 43 | 22 |
| SSc-103 | 35 | UIP | 29 | - | - |

**Table 2** Patient demographics and disease-specific data. UIP = Usual Interstitial Pneumonia. NSIP = Non-Specific Interstitial Pneumonitis. FVC = forced vital capacity, TLC = total lung capacity, DLCO = diffusion capacity. "-" data not available.

| Gene | Protein | Forward Sequence (5' to 3') | Reverse Sequence (5' to 3') |
|---|---|---|---|
| *ACTA2* | αSMA | ATCACCAACTGGGACGACAT | CATACATGGCTGGGACATTG |
| *COL1A1* | Col1a1 | CTGCTGGCAAGAGTGGTGAT | GGTGACCCTTTATGCCTCTG |
| *GAPDH* | GAPDH | CAGCCTCAAGATCATCAGCA | ACAGTCTTCTGGGTGGCAGT |
| *MAPK11* | p38β | AGTGACCAGAGGGTCAGTGC | GGGCTTGAAGCTGAGGACT |
| *MAPK12* | p38γ | GGACCAGCTGAAGGAGATCA | TTCTCCAGGAGGTTCACAGC |
| *MAPK13* | p38δ | GATGACTGGCTACGTGGTGA | CTTTCAGGATCTGGGTCAGC |
| *MAPK14* | p38α | TGCACATGCCTACTTTGCTC | CTTCTTGGTCAAGGGGTGGT |
| *PTGER1* | EP1 | GGTATCATGGTGGTGTCGTG | ATCTGGTTCCAGGAGGCAAG |
| *PTGER2* | EP2 | CCACCTCATTCTCCTGGCTA | GCCTAAGGATGGCAAAGACC |
| *PTGER3* | EP3 | CAGCTTATGGGGATCATGTG | GCTTCTCCGTGTGTGTCTTG |
| *PTGER4* | EP4 | CTCCCTGGTGGTGCTCATCT | GGTCTAGGATGGGGTTCACA |
| *PTGIR* | IP | GACCACCTGATCCTGCTGG | GCGGAAAAGGATGAAGACCC |

**Table 3** Forward and reverse human primer sequences used to determine transcript levels of the indicated gene/protein productions in qPCR experiments.
